# *Growth Factor Independent 1* is a tumor suppressor gene in colorectal cancer

**DOI:** 10.1101/370585

**Authors:** Min-Shan Chen, Yuan-Hung Lo, Xi Chen, Christopher Williams, Jessica Donnelly, Zachary Criss, Shreena Patel, Joann Butkus, Noah F. Shroyer

## Abstract

Colorectal cancer (CRC) is the third most common cancer and the third leading cause of cancer death in the United States, causing about 50,000 deaths each year. Growth Factor-Independent 1 (GFI1) is a critical zinc finger transcriptional repressor responsible for controlling secretory cell differentiation in the small intestine and colon. GFI1 plays a significant role in the development of human malignancies, including leukemia, lung cancer and prostate cancer. However, the role of GFI1 in CRC progression is largely unknown. Our results demonstrate that RNA and protein expression of *GFI1* are reduced in advanced stages of non-mucinous CRC. Subcutaneous tumor models demonstrated that the re-expression of GFI1 in 4 different human CRC cell lines inhibits tumor growth by 25-60%. To further investigate the role of Gfi1 in de novo colorectal tumorigenesis, we developed transgenic mice harboring a deletion of Gfi1 in the distal intestine driven by the CDX2cre (Gfi1^F/F^; CDX2^cre/+^) and crossed them with Apc^Min/+^ mice (Apc^Min/+^; Gfi1^F/F^; CDX2^cre/+^). Loss of Gfi1 significantly increased the total number of colorectal adenomas compared to littermate controls with an APC mutation alone. Furthermore, we found that compound (Apc^Min/+^; Gfi1^F/F^; CDX2^cre/+^) mice develop both adenomas as well as carcinoid-like tumors expressing the neuroendocrine marker chromogranin A, a feature that has not been previously described in APC-mutant tumors in mice. Collectively, these results demonstrate that Gfi1 deficiency promotes colorectal tumorigenesis, and suggest that loss of Gfi1 may promote formation of carcinoid cancers of the large intestines.

**Significance:** These findings reveal that GFI1 functions as a tumor suppressor gene in colorectal tumorigenesis.

## Introduction

Colorectal cancer (CRC) is the third most commonly diagnosed cancer in the world, with approximately 1.4 million new cases diagnosed every year. In 2016, CRC was responsible for 8% of all incident cancers and the third leading cause of cancer mortality in the United States (1). Unlike many other cancers, CRC progression is preventable and when detected early is associated with a good, long-term, prognosis (2). The 5-year survival rate is up to 90% when diagnosed with stage I or ll colorectal cancer (3,4). However, the 5-year survival rate drops to 12% for those who were diagnosed with metastatic CRC (5). The transformation from benign adenoma to malignant carcinoma is estimated to take 7-10 years (6). This gap is what allows for early detection and the prevention of death.

Transcription factors are commonly deregulated in the pathogenesis of human cancers; these transcription factors control oncogenic signaling pathways leading to tumor initiation and progression as well as metastasis. Targeting these transcription factors holds tremendous therapeutic potential for cancer treatment (7). Growth Factor-Independent 1 (GFI1) is a critical zinc finger transcriptional repressor responsible for controlling hematopoietic stem cell quiescence, myeloid and lymphoid differentiation and lymphocyte development and function (8–16). Outside of the hematopoietic system, GFI1 is required for secretory cell differentiation in the small intestine and colon as well as pulmonary neuroendocrine cell differentiation (17– 19). GFI1 regulates several transcriptional circuits whose dysregulation leads to malignancies such as leukemia, lung cancer and prostate cancer (20–23). However, whether molecular alteration of *GFI1* is associated with CRC progression is still largely unknown. Genome sequencing analysis from The Cancer Genome Atlas (TCGA) suggests that GFI1 expression is associated with less aggressive disease and better outcome. Thus, it may play a role in stalling tumor progression and metastasis (24). Here, we investigate whether dysregulation of Gfi1 expression contributes to CRC tumorigenesis. For this study, we generated GFI1 overexpression cells and knock-out mice to examine the different expression levels of GFI1 in CRC progression. Our results show that loss of *GFI1* RNA and protein in advanced stages of human CRC tissue. Re-expression of GFI1 in human CRC cell lines inhibits tumor growth in a subcutaneous tumor model. Loss of GF1I in the distal intestine accelerates the progression of malignancy in mice, promoting the formation of both high-grade adenomas and carcinoid-like tumors. Thus, our results indicate that GFI1 is a putative tumor suppressor in colorectal cancer.

## Results

### GFI1 messenger RNA and protein levels are silenced in the majority of human colon tumors

Previous studies have suggested that *GFI1* mRNA expression is associated with tumor aggressiveness in colorectal cancer. (24) To further examine the role of GFI1 in human colon tumorigenesis and metastasis, we analyzed *GFI1* mRNA expression in samples representing each stage of CRC development. We found *GFI1* expression was decreased at all cancer stages (I–IV) compared to normal colon or adenomas (normal vs all stages, p-value= 1.267e-5) (Figure 1A). Analysis of patient data showed that CRC patients with tumors that had high expression of GFI1 had a better survival rate compared with CRC patients with low expression of *GFI1* (Figure 1B). To further examine the protein expression level of GFI1 in human colorectal tumors, we analyzed an array of colonic tissue, including 145 tumors obtained from the pathology core at Baylor College of Medicine. Specificity of the GFI1 antibody was confirmed by peptide competition assay (Supplementary figure 1A). GFI1 was expressed in the nuclei of scattered cells within the normal colonic epithelium (Figure 1D, panel A, red arrows). We also observed staining of scattered cells in the lamina propria, consistent with the expression of GFI1 in bone marrow stromal cells and hematopoietic cells such as neutrophils, as previously shown (25,26). CD45 immunostaining confirmed that GFI1 positive cells in the lamina propria are hematopoietic cells (Supplementary figure 1B). We quantified the nuclear and cytoplasmic staining of GFI1 in tumor epithelium and surrounding stromal cells, and found that GFI1 was highly expressed in the tumor stroma but absent in nearly all tumor cells (Figure 1D, panel B-G). These results confirm the mRNA expression data and point to a potential role for GFI1 as a tumor suppressor in the colon. We noted that some mucinous tumors retained GFI1 expression but with higher cytoplasmic vs. nuclear staining compared to control tissue (Figure 1C, right panel and 1D, J-I), indicating that different mechanisms may control GFI1 expression in mucinous cancer tumorigenesis compared with the more common non-mucinous adenocarcinoma.

**Figure 1.**
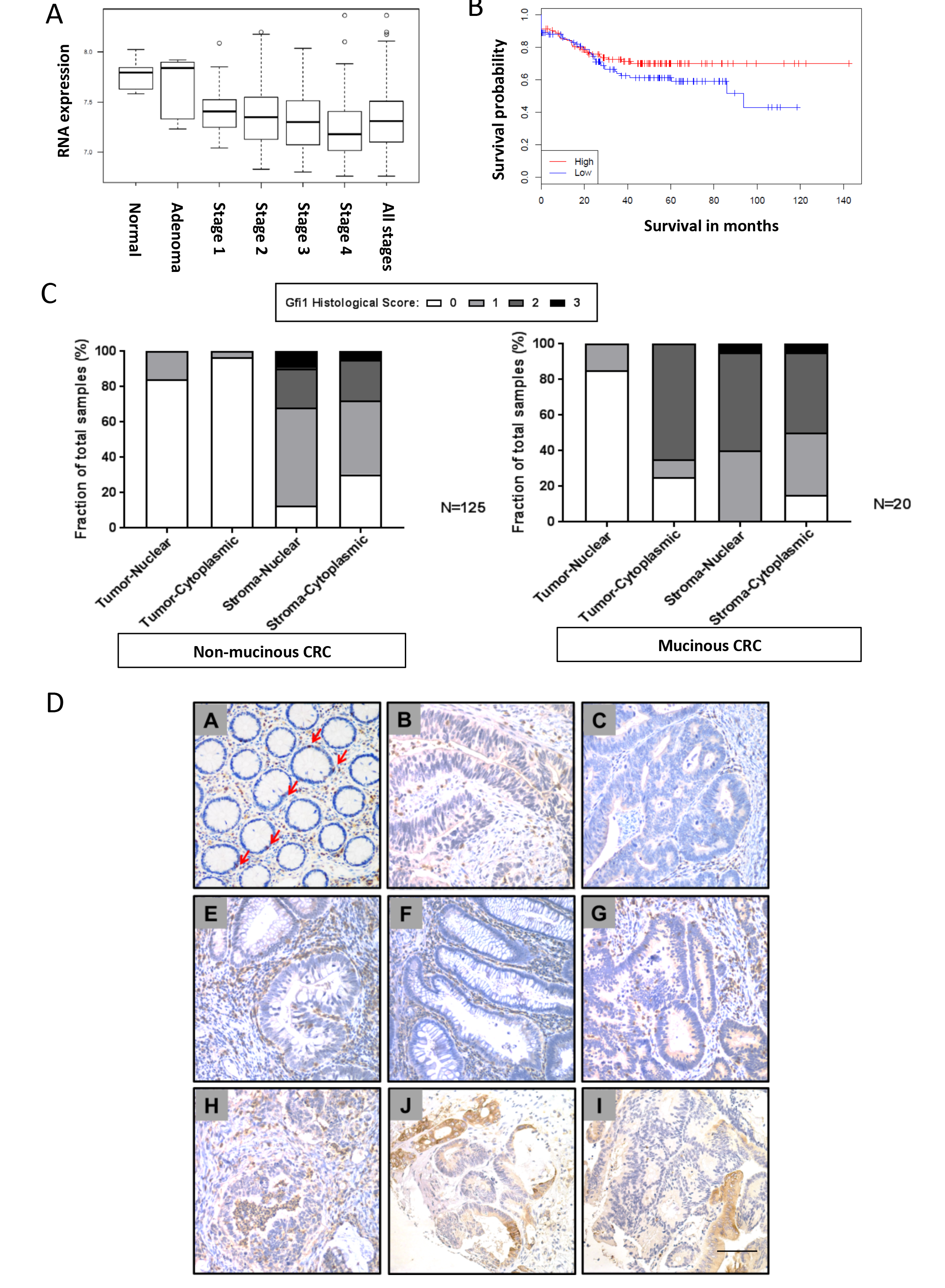
Expression of GFI1 is decreased in colorectal cancer (CRC). (A) *GFI1* expression is reduced in colorectal carcinoma compared with benign adenomas and noncancerous tissue. (B) Correlation between *GFI1* expression status and prognosis of CRC patients. (p=0.115, N=223) . (C) The distribu=on of total IHC scores for cytoplasmic or nuclear staining of Gfi1 in tumors and stroma. Top panel shows IHC scores in non-mucinous and down panel shows IHC scores in mucinous CRC. Decreased cytoplasmic and nuclear expression of GFI1 was observed in most of the CRC, whereas higher cytoplasmic Gfi1 expression was observed in mucinous CRC. Data are represented as means ± SD (D) Representative images of GFI1 staining in normal (Panel A), non-mucinous tumors (Panel B-H) and mucinous tumors (Panel J-I) are shown. Arrows indicate GFI1 positive cells in colonic epithelium. Scale bars, 100 *μm*.

### GFI1 expression has moderate effects on growth of CRC cells *in vitro*

Our results showed expression of GFI1 is reduced in CRC tumors from CRC patients (Figure 1D) and undetectable in most human CRC cell lines (Figure 2B). To investigate the effect of GFI1 re-expression on CRC progression, we generated human CRC cell lines that express GFI1 in a doxycycline-inducible manner. The human CRC cell lines, HCT116, SW480, DLD1 and LOVO were engineered to stably express doxycycline-inducible GFI1 protein, and are referred to as HCT116-hGFI1, SW480-hGFI1, DLD1-hGFI1 and LOVO-hGFI1, respectively. Control cell lines were transfected with an empty vector and were designated HCT116-control, SW480-control, DLD1-control and LOVO-control. GFI1 protein was detectable in the HCT116-hGFI1, SW480-hGFI1, DLD1-hGFI1 and LOVO-hGFI1 cells after 48 hours of induction using immunostaining (Figure 2A) and western blot (Figure 2B). Next, to examine whether or not re-expression of GFI1 affects the cell cycle, flow cytometry cell cycle analysis was performed using propidium iodide DNA staining of GFI1-expressing or control CRC cells 48 hours post induction. HCT116-hGFI1, SW480-hGFI1 showed a decrease in the number of cells in the G0/G1 phase and accumulation of cells in the S and G2/M phases. However, no significant differences were observed in any phase of the cell cycle in the DLD1-hGFI1 and LOVO-hGFI1 cell lines as compared to controls, suggesting that re-expression of GFI1 has a moderate effect on cell cycle (Figure 2C).

**Figure 2.**
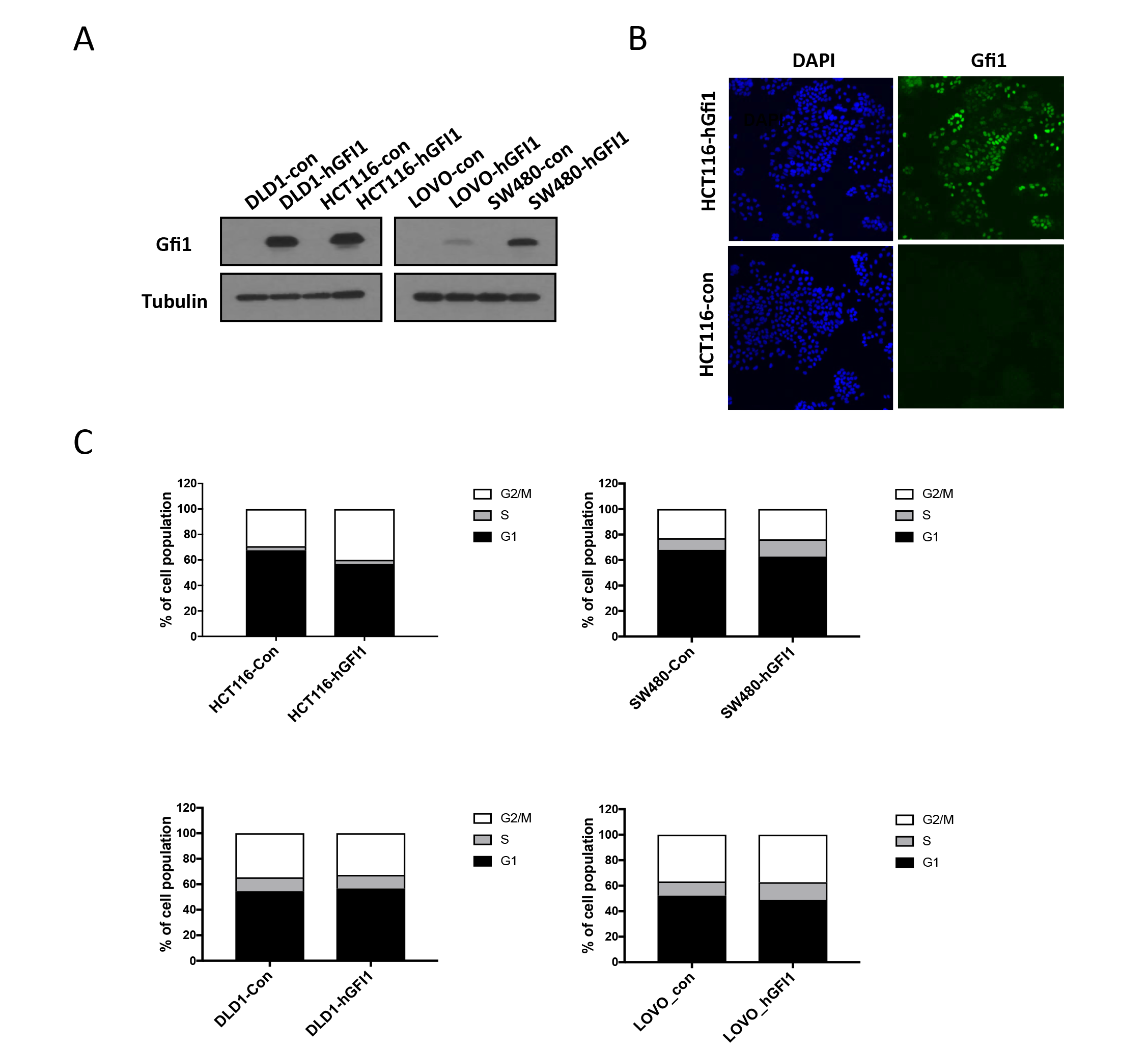
Establishment of CRC cell lines expressing doxycycline-inducible hGFI1 protein. CRC cell lines were transfected with either pINDUCER20-hGFI1 expression construct or pINDUCER20 empty vector. Cell were selected by G418 for 2 weeks and induce GFI1 expression using 1μg/mL doxycycline for 48 hours. **(A)** Western blot analysis of cell lysates was performed using anti-GFI1 antibody. (B) Cells were visualized 48 hours post induction for GFI1 expression. Scale bars, 100 *μm*. (C) Cell-cycle analysis showing the percentage of cells in G_1_, S-, and G_2_–M phases in GFI1-expressing and control CRC cells (HCA1116, SW480, DLD1 and LOVO) 48 hours post induction.

### GFI1 overexpression impairs growth of human CRC xenografts

To test whether re-expression of GFI1 inhibits cancer cell growth in vivo, we subcutaneously injected Gfi1-expressing cells (HCT116-hGFI1, SW480-hGFI1, DLD1-hGFI1 and LOVO-hGFI1) and control cells (HCT116-control, SW480-control, DLD1-control and LOVO-control) into the right and left rear flank, respectively, of immunodeficient NSG mice (Figure 3A). Mice were fed with doxycycline chow 3 days before and continuously after cancer cell transplantation to induce GFI1 expression (Figure 3B). Immunohistochemical staining showed GFI1 expression after induction by doxycycline (Figure 3G). Tumor size and body weight were measured twice a week for 4 weeks. We found that in all four CRC cell lines, GFI1 expression significantly reduced the growth of human CRC xenografts (Figure 3C-F).

**Figure 3.**
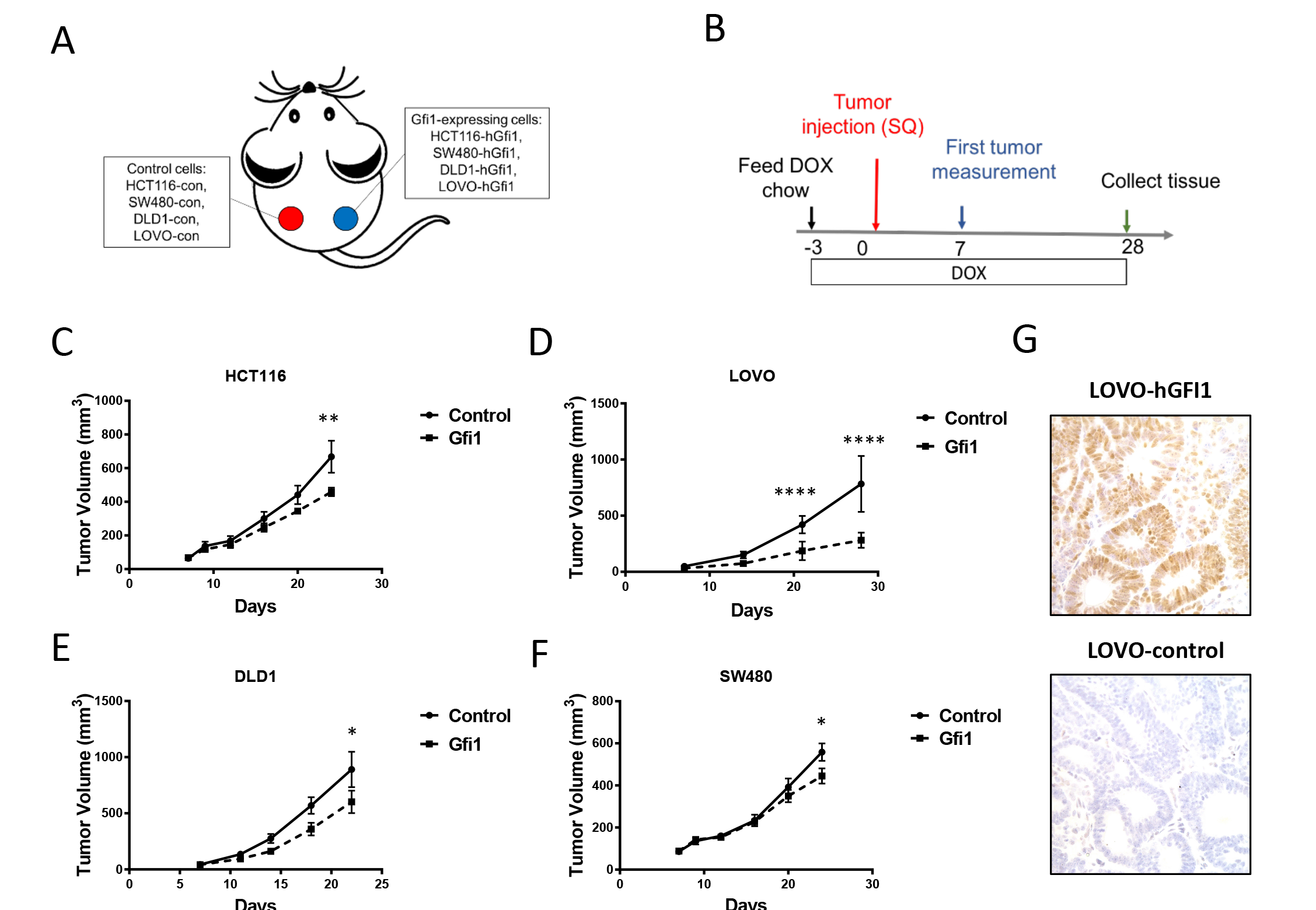
GFI1 overexpression inhibits tumor growth in subcutaneous tumor model. (A) The establishment of tumor model by subcutaneous injection. (B) Scheme of tumor injection and GFI1 induction and tissue collection. (C) Growth of subcutaneous HCT116 colon tumors (n=10). (D) Growth of subcutaneous LOVO colon tumors (n=10). (E) Growth of subcutaneous DLD1 colon tumors (n=8). (F) Growth of subcutaneous SW480 colon tumors (n=8). P value, * <0.05; *** <0.0001 (G) GFI1 expression was observed ager induction in LOVO cells by immunohistochemical staining.

### Loss of GFI1 in Distal Intestine is Associated with Colon Tumorigenesis

To investigate the effect of *Gfi1* deficiency on colorectal adenoma formation and progression, we crossed Gfi1^F/F^; CDX2^cre/+^ mice with Apc^Min/+^ mice, the best-studied model of de novo colon tumorigenesis. In adult CDX2^cre/+^ mice, Cre activity is limited to the distal intestine, which we confirmed in by crossing CDX2^cre/+^ with reporter Rosa26-tdTomato mice (Supplemental Figure 2). Apc^Min/+^; Gfi1^F/F^; CDX2^cre/+^ mice developed significantly larger and more tumors in the rectum and colon at age 4-5 months than littermates retaining Gfi1 (Apc^Min/+^; Gfi1^F/+^; CDX2^cre/+^ and Apc^Min/+^; Figure 4A-B). In contrast to control Apc^Min/+^ mice, which develop few colonic tumors, Apc^Min/+^; Gfi1^F/F^; CDX2^cre/+^ mice had a >5-fold increase in colorectal adenoma multiplicity (average number of colorectal adenomas per mouse: 29 vs. 5, Figure 4C). Colorectal adenomas from Apc^Min/+^; Gfi1^F/F^; CDX2^cre^ mice were also larger than in Apc^Min/+^ mice, with a 4-fold increase in the number of large tumors (>4 mm) and 13-fold increase in the number of medium-sized tumors (between 2-4 mm) (Figure 4D). Histological analysis confirmed the increased tumor burden, and that the increased tumor size was due to expansion in tumor cells and not large cystic structures on inflammatory infiltrate (Figures 4E; 5). PCR analysis confirmed the Cre-mediated deletion of the *Gfi1* allele (exon 4 and exon5), indicating that the larger tumors arose from Gfi1-mutant tissue (Figure 4F). Homozygous deletion of *Gfi1* in mice has been shown to affect secretory cell differentiation in the intestine with a reduced numbers of goblet and Paneth cells and the increased numbers of enteroendocrine cells. (18) We characterized mucous-secreting cells using Alcian Blue staining of acidic mucins, which are normally restricted to pre-goblet and goblet cells in the colon. In tumors from both Apc^Min/+^; Gfi1^F/F^; CDX2^cre^ and Apc^Min/+^ mice, we observed a reduction in mucous-secreting goblet cells (Figure 5A). Cells within the adenomas stained for Ki67 and PCNA, indicating that they were actively proliferating in Gfi1 knockout and control mice (Figure 5A). Consistent with their reduced secretory cell differentiation (27), Apc^Min/+^ mice developed adenomas without enteroendocrine features (Figure 5). In contrast, we found extensive expression of the enteroendocrine cell marker, Chromogranin A (CgA), in different regions of *Gfi1* knockout tumors (Figure 5A-B). CgA+ cells are scattered throughout low-grade adenomas that arose in *Gfi1* knockout mice (Figure 5A, middle panels). In high-grade adenomas with invasive features, we observed many CgA+ cells (Figure 5A, bottom panels). In some cases, these CgA+ cells appeared to have delaminated from the glandular structures of the tumor. Interestingly, we also found carcinoid-like cells which clustered together in the stroma surrounding hyperplastic crypt epithelium in the colons of Apc^Min/+^; Gfi1^F/F^; CDX2^cre^ mice. (Figure 5B) These carcinoid-like cell clusters generally did not express KI67 nor PCNA, suggesting that they are well-differentiated and slow-growing tumors. RT-qPCR analysis showed increased expression of chromogranin A and decreased expression of the goblet cell marker, *Muc2*, and no significant differences in Wnt target genes and stem cell genes in Gfi1 knockout tumors compared to control tumors. (Supplementary Figure 3). Taken together, these results indicated that *Gfi1* deficiency greatly promotes tumorigenesis and formation of carcinoid-like tumors in the colon and rectum.

**Figure 4.**
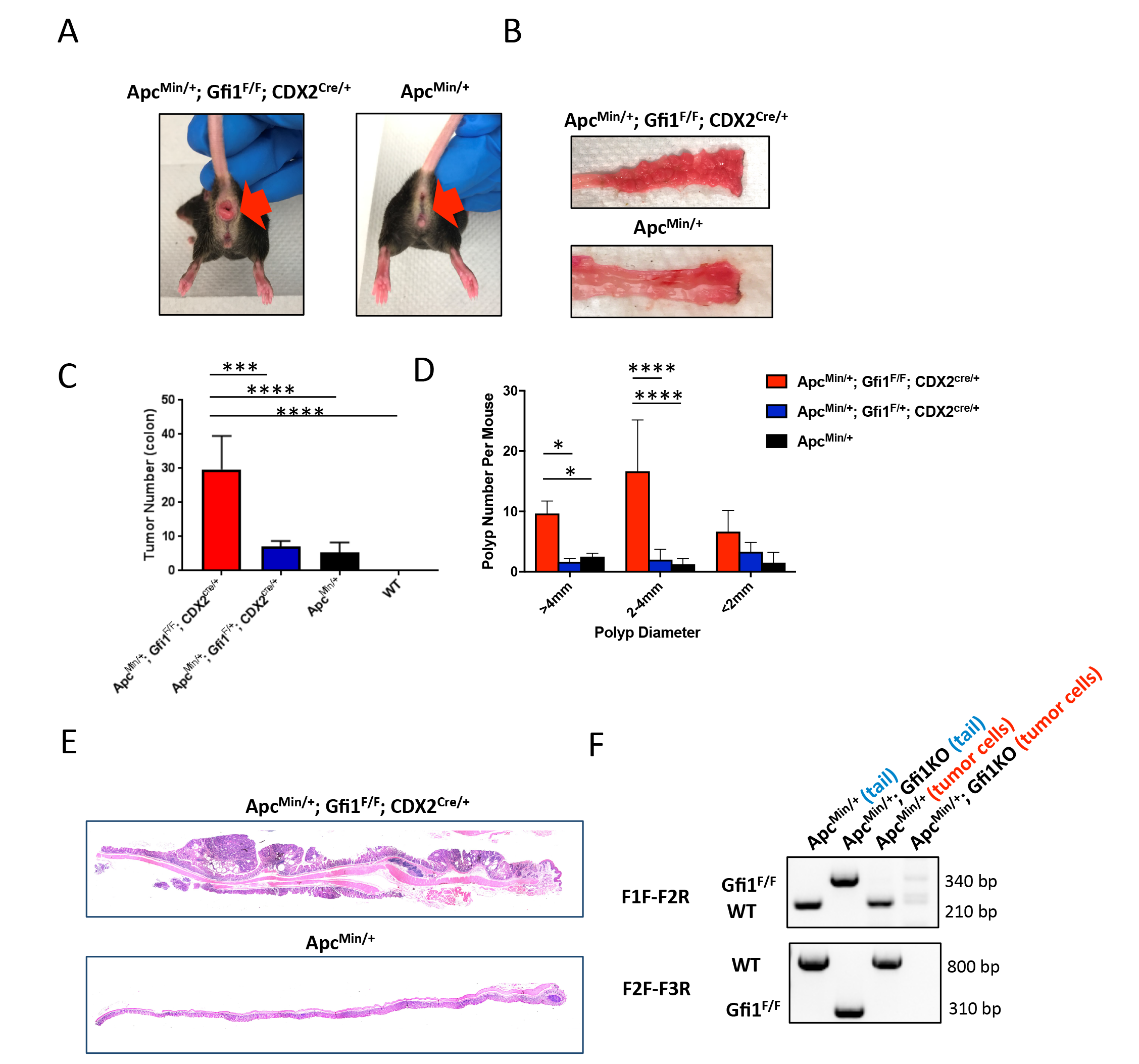
GFI1 loss leads to increased tumor burden in the colon of Apc^Min/+^ mice. Representative photographs of rectum (A) and longitudinally opened distal colon (B) from Apc^Min/+^; Gfi1^F/F^; CDX2^Cre/+^ and Apc^Min/+^; Gfi1^wt^; CDX2^wt^ (Apc^Min/+^). Analysis of polys in the colon, (C) number of polyps per mouse; (D) diameter of polyps in Apc^Min/+^; Gfi1^F/F^; CDX2^Cre/+^ (n=3), Apc^Min/+^; Apc^Min/+^; Gfi1^F/+^; CDX2^Cre/+^ (n=3), Apc^Min/+^ (n=5), wildtype, WT (n=5). Error bars represent SD. (E) Representa=ve histopathology of tumors from Apc^Min/+^; Gfi1^F/F^; CDX2^Cre/+^ and Apc^Min/+^ mouse. (F) A PCR-based assay iden=fied the Gfi1^F/F^ and Gfi1wt alleles in mouse tails and tumor organoids from Apc^Min/+^; Gfi1^F/F^; CDX2^Cre/+^ and Apc^Min/+^ mouse. P value, * <0.05; **** <0.0001

**Figure 5.**
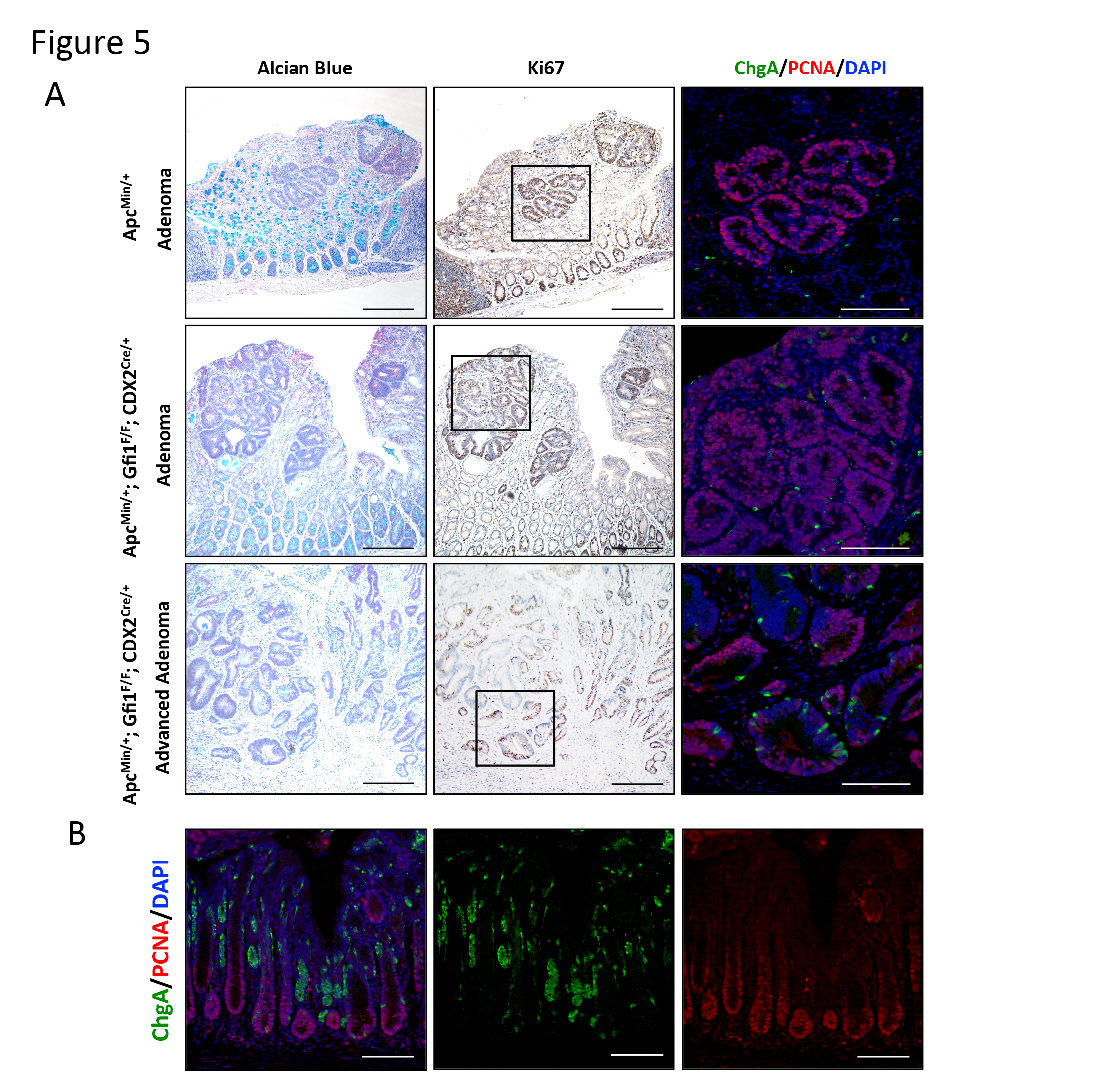
Histological analysis of GFI1 knockout tumors. Control (Apc^Min/+^) and GFI1 knockout tumors (Apc^Min/+^; Gfi1^F/F^; CDX2^Cre/+^) were analyzed by histology. (A) To evaluate mucous-secreting goblet cells in these tumors, Alcian blue staining was used to visualize mucous globules. GFI1 knockout mice showed a dramatic decrease in the number of goblet cells in the colon. Proliferating cells were analyzed by Ki67 or PCNA staining and enteroendocrine cells were analyzed by ChgA staining. Fluorescent images on the right panels showed enteroendocrine cell numbers were markedly increased in advanced adenomas and slightly increased in adenoma in GFI1 knockout mice. Leg panels: hematoxylin & Eosin and Alcian blue staining, scale bar: 200 *μ*m. Middle panels: KI67 and hematoxylin staining, scale bar: 200 *μ*m. Right panels: chromogranin A (ChgA), PCNA and nuclei staining (DAPI), scale bar: 100 *μ*m. (B) Carcinoid tumors arose from enteroendocrine cells were observed in hyperplasia area in GFI1 knockout tumors. In the hypoplasia area of GFI1 knockout colon, small clusters of enteroendocrine cells (green) scanered throughout mucosal layer. Scale bars: 100 μm.

## Materials and Methods

### Cell culture

The human colorectal cancer cell line HCT116, SW480, DLD1 and LOVO, as well as the human embryonic kidney cell lines HEK-293T were purchased from American Type Culture Collection (ATCC). All cells were grown in RPMI (Corning, 10-040-CV) supplemented with 10% fetal bovine serum (Bioexpress, S1200-500), penicillin and streptomycin (Lonza, 17-602E). The doxycycline inducible pINDUCER HCT116, SW480, DLD1 and LOVO cell lines were maintained in the same condition with tetracycline-free fetal bovine serum (Corning, #35-075-CV) and periodically selected by supplemental 1μg/ml G418 (Sigma, #A1720).

### Generation of doxycycline-inducible cell lines

The pENTR-GFI1 plasmid (Plasmid #16168) was obtained from Addgene and cloned into pINDUCER20-Neo (28) using the Gateway Cloning Technology (Invitrogen) as previously described (29). The resulting pINDUCER20-GFI1-Neo plasmid was verified by DNA sequencing (BCM, sequence core). Virus was generated by transfection of 293T cells grown to 70% confluence in a 6-cm tissue culture plate with 2.25 μg envelope plasmid, pCMV-VSV-G (Addgene #8454) and 0.25 μg packaging plasmid, psPAX2 (Addgene #12260) and 2.5 μg pINDUCER20-GFI1-Neo plasmid or control plasmid pINDUCER20-Neo using Lipofectamine 2000 (Invitrogen) following the manufacturer’s transfection protocol. Eighteen hours after transfection, the media in each well was replaced with fresh media. After 48 hours of viral production, virus-containing media was collected, pooled, filtered through a 0.45-μm polyvinylidene difluoride membrane (Millex), and added to human colorectal cancer cell lines at 60% confluence in 10-cm tissue culture dishes (Corning). Infection was allowed to proceed for overnight, and then the media weas replaced. After a 72-hour recovery period, infected cells were selected with 1 μg/mL G418 (Sigma, #A1720).

### Western blot

Cell lysates of the treated cells were isolated by incubation with RIPA buffer, containing 150 mM NaCl, 1 mM EDTA pH 8.0, 25mM Tris, 0.1% SDS, 0.5% sodium deoxycholate, 1% NP-40, phosphatase inhibitor cocktails (Sigma, P5726 and P0044) and the protease inhibitors cocktail (Calbiochem, #539134). Cell lysates were separated on acrylamide gels, transferred to a PVDF membrane (Bio-Rad), and probed with the Gfi1 antibody (1:1000 Santa Cruz, sc8558) and tubulin antibody (DSHB, 6G7). Bands were visualized by a chemiluminescence-based detection method (Fisher/Pierce) that used horseradish peroxidase-conjugated secondary antibodies (Cell Signaling, #7076 and Santa Cruz, sc-2056).

### Mice

NSG mice (#00557) were obtained from The Jackson Laboratory. For xenograft studies, male NSG mice were randomly divided into experimental groups. Mice were subcutaneously injected with 2 × 10^6^ HCT116, SW480, DLD1 or LOVO cells in a 100 μl medium (50% matrigel in serum-free RPMI1640). To induce Gfi1 expression, mice were fed doxycycline diet (625 mg/kg, Envigo Teklad) 3 days before inoculation of HCT116, SW480, DLD1 or LOVO cells. Tumor volume (mm^3^) was calculated using the following formula: tumor volume (mm^3^) = length (mm) × width^2^ (mm^2^) × 0.5. Apc^Min/+^ mice were purchased from the Jackson Laboratory (#002020). CDX2^cre/+^ mice (30) were obtained from Dr. Jason Heaney at Baylor College of Medicine. Gfi1^F/F^ mice (12) were obtained from Dr. Jinfang Zhu at National Institutes of Health. All mouse studies were approved by Institutional Animal Care and Use Committee (IACUC).

### Analysis of cell cycle by flow cytometry

GFI1-expressing cells and control cells were treated with 1 μg/mL doxycycline for 48 hours. Cells were trypsinized and fixed in cold 70% ethanol for 10 minutes and then stained with propidium iodide (PI) solution (1 μg/μL PI and 0.125% RNaseA; Sigma) at room temperature for 15 minutes. Approximately 10,000/sample cells were analyzed using LSRFortessa (Becton Dickinson).

### Tissue staining

Colon and tumors were fixed with 4% PFA at 4 °C overnight, washed with PBS and then transferred to 70% ethanol for paraffin-embedding. Tissues were sectioned at 5 μm thickness. Paraffin-embedded sections were deparaffinized and rehydrated before staining. For immunohistochemistry, antigen retrieval was achieved in sodium citrate buffer (10 mM sodium citrate pH 6.0). Slides were incubated in 3% H_2_O_2_ solution in methanol at room temperature for 10 minutes to block endogenous peroxidase activity. After washing, slides were blocked in Avidin/Biotin solution (Vector Laboratories, SP2001) for 30 minutes and then in 5% normal goat or donkey serum in PBST at room temperature for 1 hour. After blocking, slides were incubated with primary antibody at 4°C overnight. Slides were washed by PBST at room temperature for 5 minutes three times and incubated with secondary antibody at room temperature for 0.5 hour. Slides were washed by PBST and then ABC reagent applied (Vector Laboratories, PK-6101) for 0.5 hour. Slides were washed by PBST and signals were detected by DAB (Vector Laboratories, SK-4100). The following antibodies were used: goat Gfi1 (1:100 Santa Cruz, sc-8558), rabbit Ki67 (1:1000 Leica, NCL-Ki67p), biotinylated rabbit anti-goat IgG antibody (Vector Laboratories, BA-5000), biotinylated goat anti-rabbit IgG antibody (Vector Laboratories, BA-1000). For immunofluorescent staining, deparaffinization and rehydration procedures were described above. Slides were blocked with 5% serum at room temperature for 1 hour or M.O.M. reagent if a mouse antibody was used (Vector Laboratories, PK-2200). Primary antibody was diluted in 5% serum in PBST and incubated with tissues at 4 °C for overnight. Slides were washed by PBST (at room temperature for 3 minutes for 3 times) and incubated with secondary antibody at room temperature for 1 hour. After 3 washes, slides were stained with DAPI (Thermo/Fisher, #62248) at room temperature for 15 minutes. Slides were washed twice with PBS. Slides were mounted with mounting solution (Vector Laboratories, H-1000). The following antibodies were used: mouse PCNA (1:1000, Cell Signaling, #2586), mouse E-cadherin (1:250 Abcam, ab76055), rabbit beta-catenin (1:500 Santa Cruz, sc-7199), mouse CD45 (1:50, DSHB, H5A5-s), Chromogranin A (1:1000, immunostar, 20085). For goblet cell staining, slides were stained with Alcian blue (Thermo/Fisher, 88043) and followed manufacturer’s protocol.

### Tissue array analysis

Colon cancer tissue microarray was obtained from the Pathology Core at Baylor College of Medicine. The extent of nuclear and cytoplasmic Gfi1 expression in the epithelium of each tissue section was assessed and scored with numeric numbers; for extent, such that: 0, no staining; 1, 1%–25%, 2, 26%–50%, 3, 51%–75% and 4, 76%-100% of epithelial cells in each tissue section.

### RNA analysis

To determine if GFI1 expression was significantly different between normal adjacent colon, adenocarcinoma, and colorectal cancer (CRC), we examined the expression levels between normal adjacent colon, adenocarcinoma and each stage. Vanderbilt Medical Center/Moffitt Cancer Center colon cancer microarray data set (n=250 cancers, n=10 normal adjacent colon samples and n=6 adenocarcinoma) was downloaded from Gene Expression Omnibus (GSE17538). The microarray data were normalized using Robust Multichip Averaging (RMA) algorithm implemented in Bioconductor package Affy. Log2 transformed data were used for downstream analyses. The Wilcoxon and Kruskal-Wallis rank sum tests were used to test for significance between each stage and the normal adjacent tissue.

### Survival analysis

CRC patients from VMC/MCC with GFI1 high expression were defined as having greater than median expression of GFI1 and were compared to the low expression group (less than the median expression value). Kaplan-Meier analysis was performed, comparing patients with high GFI1L expression (red line) to low GFI1 expression (blue line). While not achieving statistical significance, patients with high GFI1 expression had better relapse-free survival than patients with low GFI1 expression (Log-rank test p=0.1692).

## Discussion

Colorectal cancer (CRC) is a highly heterogeneous disease caused by different molecular pathogenic pathways with diverse clinical outcomes, response to cancer therapy and morphological features. The dysregulation of several transcription factors is known to contribute to intestinal tumorigenesis (31). Growth factor independent 1 (GFI1) is a zinc finger transcriptional repressor implicated in diverse developmental contexts. (16,32,33) Recent studies indicated that dysregulation of zinc finger containing proteins (ZNFs) contributes to tumorigenesis in different aspects. Several studies have suggested that ZNFs act as double-edged sword during tumorigenesis in different cancer types. (34) GFI1 is critical to several transcriptional circuits whose dysregulation results in oncogenesis in the hematopoietic system, lung cancer, prostate cancer and medulloblastoma. (20–22,35,36) RNA expression in CRC tumors based on tumor stage, lymph node status, distant metastasis and vascular invasion at the time of surgery has shown that higher expression of GFI1 was associated with less tumor aggressiveness. (24) Previous studies showed GFI1 downregulation promotes inflammation-induced CRC metastasis through the activation of inflammatory signaling that enhances the invasive behavior of CRC cells. (37) However, the role of GFI1 in CRC initiation and progression is still largely unknown. In this study, we found the expression of GFI1 is progressively lost in human CRC tissues. Moreover, we found that CRC patients with lower GFI1-expressing tumors had a poor survival rate. Consistent with these findings, in human colon cancer cell lines, expression of Gfi1 inhibits cancer cell growth in a subcutaneous tumor model *in vivo*. Most importantly, we showed that deletion of *Gfi1* in the intestinal epithelium in a beta-catenin-driven transgenic mouse model of tumorigenesis results in an increased colonic tumor burden and size. As previous studies have shown that *GFI1* is a negative regulator of CRC metastasis and that downregulation of *GFI1* enhances cell migration and invasion (37), our study supports GFI1’s function as a tumor suppressor in CRC.

The molecular mechanisms that result in loss of GFI1 protein in advanced CRC (Figure 1) are not well understood, and may occur by multiple mechanisms. *GFI1* is not subject to frequent point mutation or copy number variation as reported in the COSMIC or ONCOMINE databases. Instead, it is likely that GFI1 expression is regulated by epigenetic and mechanisms, such as DNA methylation or chromatin remodeling. Previous studies suggested that the expression of *Gfi1* could be regulated through a polycomb repressive complex 2 (PRC2)-dependent mechanism. PRC2 is an important chromatin modifier involved in the maintenance of transcriptional silencing by inducing mono, di- and trimethylation of histone 3 at lysine 27 (H3K27me3). (38) EZH2, a member of the PRC2 genes, is up-regulated and correlated with poor prognosis in human CRCs. (39,40) In medulloblastoma, ChIP-seq analysis showed that EZH2 transcriptionally suppresses *Gfi1* directly, resulting in the inhibition of tumor growth (36) In addition, the expression of Gfi1 could be also regulated by the components within the tumor microenvironment. Previous studies suggested that immune cell cytokines, such as IL1α, TGFβ or IFNγ could contribute to Gfi1 deregulation, thereby changing cancer cell behaviors.(37) Finally, in the intestinal epithelium, our previous studies showed that Gfi1 is a direct target of Atoh1, a master transcription factor that controls intestinal secretory cell fate determination. (18,41) In CRCs, Atoh1 functions as a CRC tumor suppressor (42). The expression of Atoh1 is silenced due to either CpG island hypermethylation or deletion. Therefore, these results indicated that downregulation of Gfi1 might be directly regulated to a loss of Atoh1 in CRCs, leading to transformation of an early adenoma into carcinomas and metastatic tumors (42). Future studies are needed to determine the mechanisms that control GFI1 stability and co-factor interactions in CRC tumorigenesis.

Mucinous CRC represents a distinct clinical and histopathological subtype of CRC, and is found in 10–15% of patients with CRC. Mucinous CRC is associated with rapid disease progression and poor prognosis. It has been shown that mucinous CRC has different molecular alternations and genetic subtypes, when compared to non-mucinous CRC (43). Mucinous CRC shows more frequent microsatellite instability and oncogenic KRAS, BRAF and PIK3CA mutations than non-mucinous CRC (44,45). In this study, we found that *GFI1* expression is nearly absent in non-mucinous colorectal adenocarcinoma, but highly expressed in mucinous CRC (Figure 1), suggesting that GFI1 might have distinct functions in mucinous CRC as compared with non-mucinous colorectal adenomas and adenocarcinomas.

Carcinoids/gastrointestinal neuroendocrine tumors are composed of hormone-producing endocrine-like cells (46). These neuroendocrine tumors often release hormones into the bloodstream where they affect the function of other parts of the digestive system and a variety of other organ processes. A previous study suggested that overexpression of gastrointestinal hormones (gastrin) can drive formation of gastrointestinal tumors (47). It is possible that overexpression of hormones from carcinoid-like tumors in *Gfi1* knockout mice enhanced the formation and progression of colorectal adenomas in the context of *Apc* mutation. Canonical Wnt/β-catenin signaling is essential for the maintenance of intestinal stem cells and secretory cell differentiation (48,49). Within secretory progenitors, GFI1 promotes goblet and Paneth cell fate by repressing *Neurog3* expression [Shroyer et al, 2005; Bjerknes et al, 2010] . Previous work indicated that Neurog3+ early precursors of enteroendocrine cells can respond to hyperactivation of β-catenin by developing serotonin-expressing adenomas in the small intestine (50). It is likely that in our mice, deleting *Gfi1* in the intestinal epithelium resulted in more Neurog3+ enteroendocrine-cell progenitors, and subsequent loss of heterozygosity of *APC* within this expanded pool of Neurog3+ progenitors lead to endocrine-like tumors. Our finding further suggests that the biological basis for colorectal neuroendocrine tumors (NETs) may be distinct from NETs found in other tissues such as small intestine, stomach, and pancreas. This hypothesis is further supported by the observation in the COSMIC database that colorectal NETs primarily bear mutations in *p53, APC,* and *β-catenin*, whereas other gastroenteropancreatic NETs bear a different mutation profile including alterations of *MEN1*, *DAXX*, and *ATRX.*

In summary, genetic knockouts in a transgenic mouse model and molecular analyses in human CRC samples support the role of GFI1 as a tumor suppressor. Furthermore, our findings suggest that GFI1 may have distinct roles in different histological subtypes of CRC. Therefore, elucidating the basic mechanisms of GFI1 function, its target genes and interacting proteins as well as identification of pathways that control *GFI1* expression may help to define rationally targeted therapies for mucinous, carcinoid, and non-mucinous colorectal cancers.

## Acknowledgments

We thank the Texas Medical Center Digestive Disease Center for their support with funding from the National Institutes of Health (P30DK56338), the Intestinal Stem Cell Consortium with funding from the NIH through grants (U01 DK103168, U01 DK103168-03S), NIH grants R01 CA142826 (N.F.S.) and F99 CA212433 (Y.H.L). We thank the Integrated Microscopy core at Baylor College of Medicine for their support with funding from the NIH (DK56338 and CA125123), the Cytometry and Cell Sorting Core at Baylor College of Medicine with funding from the National Institutes of Health (P30 AI036211, P30 CA125123, and S10 RR024574), the Pathology and Histology Core (HTAP) at Baylor College of Medicine with grant from NCI (NCI-CA125123) and the Dan L. Duncan Cancer Center.

---

**Supplementary figure 1.**
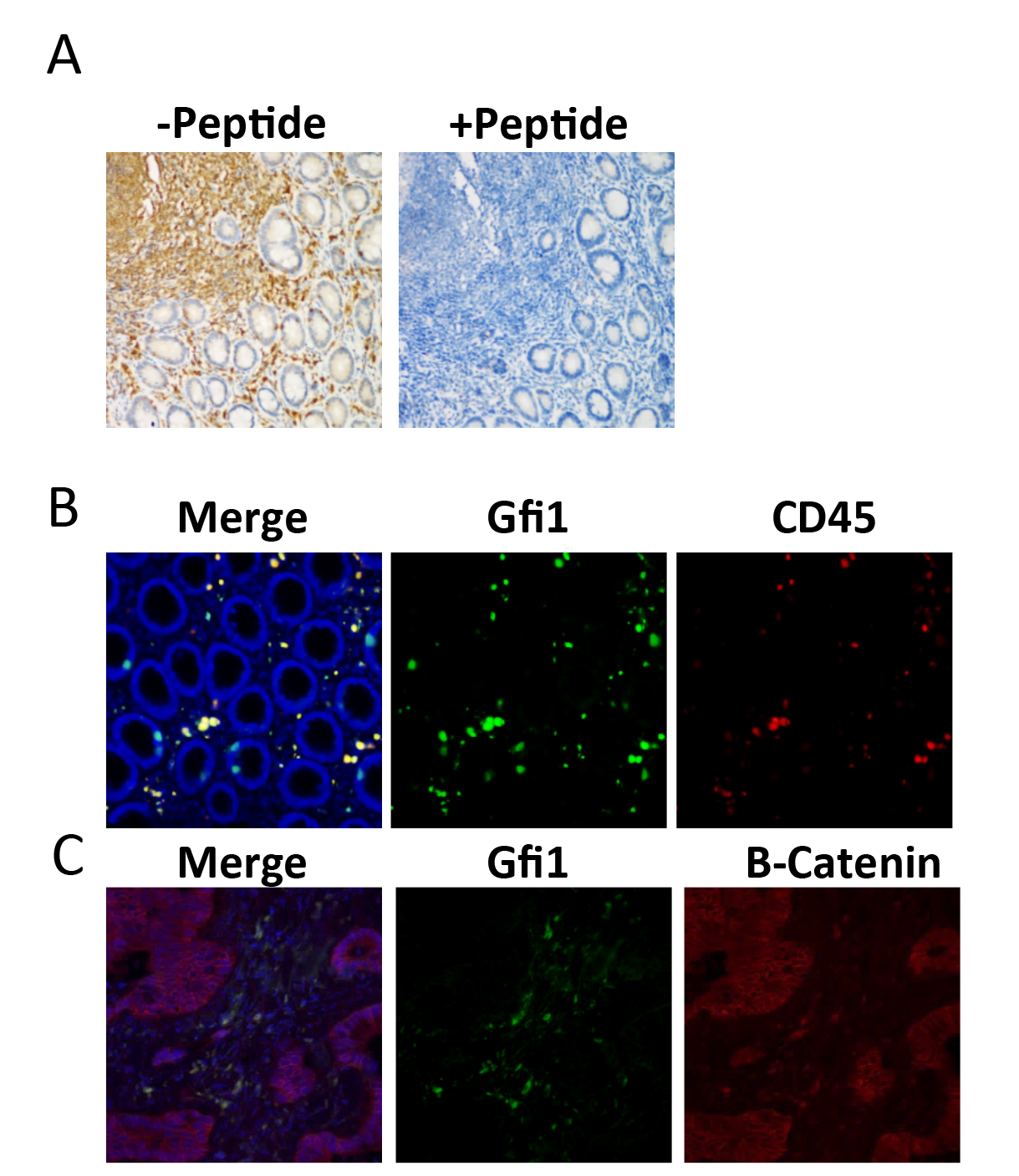
(A) Immunohistochemical analysis of tumor-adjacent normal tissue using Gfi1 antibody (Santa cruz, sc-8558) with blocking pep=de (right) or without peptide (leg). (B) Immunofluorescent staining of Gfi1 and CD45 staining shows that immune cells outside of colonic epithelium are Gfi1 and CD45 positive cells. Blue: DAPI, green: Gfi1, red: CD45. Immunofluorescent staining of Gfi1 and Β-Catenin staining shows that the expression of Gfi1 is lost in cancer cells.

**Supplementary figure 2.**
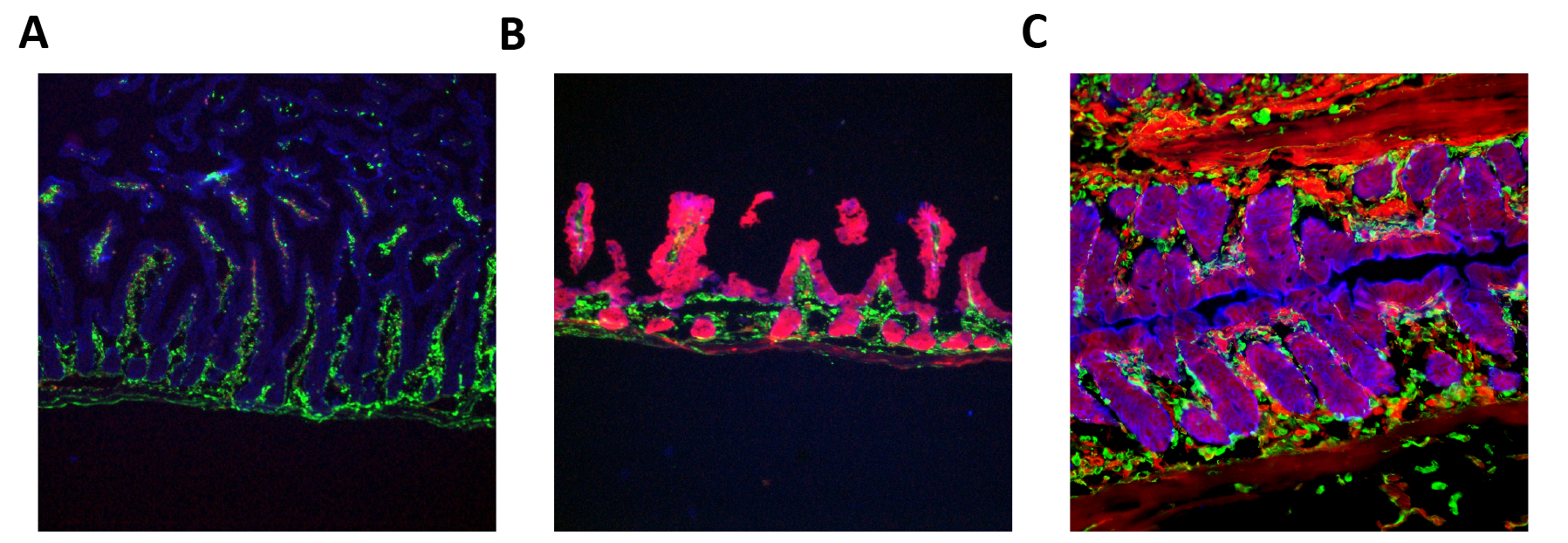
Intestinal tissues collected from CDX2^cre/+^; Rosa26-tdTomato mouse showed CDX2c^re/+^ expression (red) in proximal small intestine (A), distal small intestine (B) and colon (C). Green: Vimentin; Blue: DAPI.

**Supplementary figure 3.**
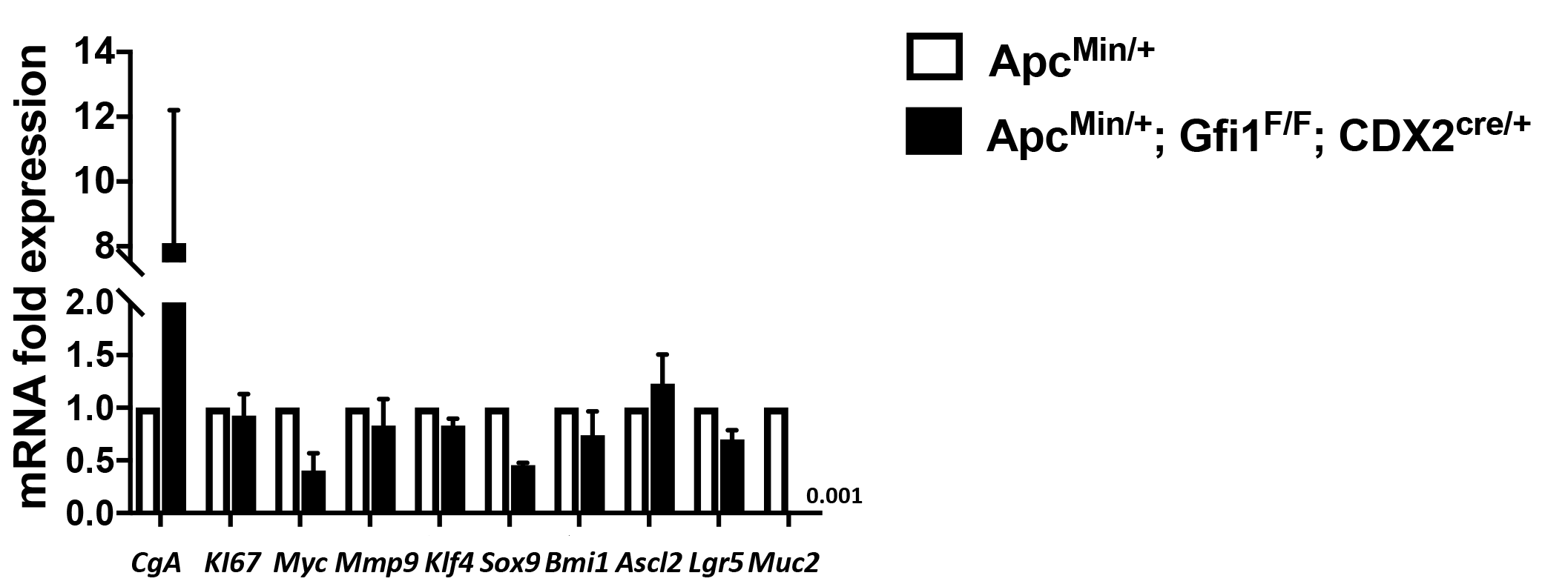
(A-B) Enteroendocrine cell marker (*ChgA*), proliferating cell marker (*Ki67*), WNT signaling markers (*Myc,Mmp9*), stem cell marker (S*ox*9, B*mi*1, A*scl*2, L*gr*5) and goblet cell marker (*Muc2*) were analyzed using qPCR .

## References

1. Siegel RL, Miller KD, Jemal A. Cancer statistics, 2016. CA Cancer J Clin [Internet]. 2016;66:7–30. Available from: http://www.ncbi.nlm.nih.gov/pubmed/26742998%5Cnhttp://onlinelibrary.wiley.com/store/10.3322/caac.21332/asset/caac21332.pdf?v=1&t=io59driw&s=f5d356a70462e7927a65b0ec8811c011b92d7469

2. Gonzalez-Pons M, Cruz-Correa M. Colorectal Cancer Biomarkers: Where Are We Now? Biomed Res Int [Internet]. 2015;2015:149014. Available from: http://www.ncbi.nlm.nih.gov/pmc/articles/PMC4461726/?tool=pubmed%5Cnhttp://dx.doi.org/10.1155/2015/149014

3. Grützmann R, Molnar B, Pilarsky C, Habermann JK, Schlag PM, Saeger HD, et al. Sensitive detection of colorectal cancer in peripheral blood by septin 9 DNA methylation assay. PLoS One. 2008;3.

4. Losso GM, Moraes RDS, Gentili AC, Messias-Reason IT. Microsatellite instability--MSI markers (BAT26, BAT25, D2S123, D5S346, D17S250) in rectal cancer. Arq Bras Cir Dig [Internet]. 2012;25:240–4. Available from: http://www.ncbi.nlm.nih.gov/pubmed/23411922

5. Bupathi M, Wu C. Biomarkers for immune therapy in colorectal cancer: Mismatchrepair deficiency and others. J. Gastrointest. Oncol. 2016. page 713–20.

6. Fichera A. Risk of progression of advanced adenomas to colorectal cancer by age and sex: Estimates based on 840,149 screening colonoscopies - Abstractor’s comment. Dis. Colon Rectum. 2008. page 486–7.

7. Bhagwat AS, Vakoc CR. Targeting Transcription Factors in Cancer. Trends in Cancer. 2015. page 53–65.

8. Karsunky H, Zeng H, Schmidt T, Zevnik B, Kluge R, Schmid KW, et al. Inflammatory reactions and severe neutropenia in mice lacking the transcriptional repressor GFi1. Nat Genet. 2002;30:295–300.

9. Person RE, Li FQ, Duan Z, Benson KF, Wechsler J, Papadaki HA, et al. Mutations in proto-oncogene GFI1 cause human neutropenia and target ELA2. Nat Genet. 2003;34:308–12.

10. Zeng H, Yücel R, Kosan C, Klein-Hitpass L, Möröy T. Transcription factor Gfi1 regulates self-renewal and engraftment of hematopoietic stem cells. EMBO J. 2004;23:4116–25.

11. Hock H, Hamblen MJ, Rooke HM, Schindler JW, Saleque S, Fujiwara Y, et al. Gfi-1 restricts proliferation and preserves functional integrity of haematopoietic stem cells. Nature. 2004;431:1002–7.

12. Zhu J, Jankovic D, Grinberg A, Guo L, Paul WE. Gfi-1 plays an important role in IL-2-mediated Th2 cell expansion. Proc Natl Acad Sci U S A [Internet]. 2006;103:18214–9. Available from: http://www.pubmedcentral.nih.gov/articlerender.fcgi?artid=1654136&tool=pmcentrez&rendertype=abstract

13. Rathinam C, Klein C. Trancriptional repressor Gfi1 integrates cytokine-receptor signals controlling B-cell differentiation. PLoS One. 2007;2.

14. Pargmann D, Yücel R, Kosan C, Saba I, Klein-Hitpass L, Schimmer S, et al. Differential impact of the transcriptional repressor Gfi1 on mature CD4+ and CD8+ T lymphocyte function. Eur J Immunol. 2007;37:3551–63.

15. Zarebski A, Velu CS, Baktula AM, Bourdeau T, Horman SR, Basu S, et al. Mutations in Growth Factor Independent-1 Associated with Human Neutropenia Block Murine Granulopoiesis through Colony Stimulating Factor-1. Immunity. 2008;28:370–80.

16. Hock H, Orkin SH. Zinc-finger transcription factor Gfi-1: versatile regulator of lymphocytes, neutrophils and hematopoietic stem cells. Curr Opin Hematol [Internet]. 2006;13:1–6. Available from: http://www.ncbi.nlm.nih.gov/pubmed/16319680

17. Bjerknes M, Cheng H. Cell Lineage metastability in Gfi1-deficient mouse intestinal epithelium. Dev Biol. 2010;345:49–63.

18. Shroyer NF, Wallis D, Venken KJT, Bellen HJ, Zoghbi HY. Gfi1 functions downstream of Math1 to control intestinal secretory cell subtype allocation and differentiation. Genes Dev. 2005;19:2412–7.

19. Linnoila RI, Jensen-Taubman S, Kazanjian A, Grimes HL. Loss of GFI1 impairs pulmonary neuroendorine cell proliferation, but the neuroendocrine phenotype has limited impact on post-naphthalene airway repair. Lab Investig. 2007;87:336–44.

20. Kazanjian A, Wallis D, Au N, Nigam R, Venken KJT, Cagle PT, et al. Growth factor independence-1 is expressed in primary human neuroendocrine lung carcinomas and mediates the differentiation of murine pulmonary neuroendocrine cells. Cancer Res. 2004;64:6874–82.

21. Dwivedi PP, Anderson PH, Tilley WD, May BK, Morris HA. Role of oncoprotein Growth Factor Independent-1 (GFI1) in repression of 25-hydroxyvitamin D 1alpha-hydroxylase (CYP27B1): A comparative analysis in human prostate cancer and kidney cells. J Steroid Biochem Mol Biol. 2007;103:742–6.

22. Dwivedi PP, Anderson PH, Omdahl JL, Grimes HL, Morris HA, May BK. Identification of growth factor independent-1 (GFI1) as a repressor of 25-hydroxyvitamin D 1-alpha hydroxylase (CYP27B1) gene expression in human prostate cancer cells. Endocr Relat Cancer. 2005;12:351–65.

23. Huang M, Hu Z, Chang W, Ou D, Zhou J, Zhang Y. The growth factor independence-1 (Gfi1) is overexpressed in chronic myelogenous Leukemia. Acta Haematol. 2009;123:1–5.

24. Cancer Genom Atlas. Comprehensive molecular characterization of human colon and rectal cancer. Nature [Internet]. 2012;487:330–7. Available from: http://www.pubmedcentral.nih.gov/articlerender.fcgi?artid=3401966&tool=pmcentrez&rendertype=abstract

25. D’Souza S, Del Prete D, Jin S, Sun Q, Huston AJ, Kostov FE, et al. Gfi1 expressed in bone marrow stromal cells is a novel osteoblast suppressor in patients with multiple myeloma bone disease. Blood. 2011;118:6871–80.

26. Hock H, Hamblen MJ, Rooke HM, Traver D, Bronson RT, Cameron S, et al. Intrinsic requirement for zinc finger transcription factor Gfi-1 in neutrophil differentiation. Immunity. 2003;18:109–20.

27. Ueo T, Imayoshi I, Kobayashi T, Ohtsuka T, Seno H, Nakase H, et al. The role of Hes genes in intestinal development, homeostasis and tumor formation. Development [Internet]. 2012;139:1071–82. Available from: http://dev.biologists.org/cgi/doi/10.1242/dev.069070

28. Meerbrey KL, Hu G, Kessler JD, Roarty K, Li MZ, Fang JE, et al. The pINDUCER lentiviral toolkit for inducible RNA interference in vitro and in vivo. Proc Natl Acad Sci [Internet]. 2011;108:3665–70. Available from: http://www.pnas.org/cgi/doi/10.1073/pnas.1019736108

29. Katzen F. Gateway ^®^ recombinational cloning: a biological operating system. Expert Opin Drug Discov. 2007;2:571–89.

30. Hinoi T, Akyol A, Theisen BK, Ferguson DO, Greenson JK, Williams BO, et al. Mouse model of colonic adenoma-carcinoma progression based on somatic Apc inactivation. Cancer Res. 2007;67:9721–30.

31. Lee TI, Young RA. Transcriptional regulation and its misregulation in disease. Cell. 2013. page 1237–51.

32. Grimes HL, Chan TO, Zweidler-McKay P a, Tong B, Tsichlis PN. The Gfi-1 proto-oncoprotein contains a novel transcriptional repressor domain, SNAG, and inhibits G1 arrest induced by interleukin-2 withdrawal. Mol Cell Biol [Internet]. 1996;16:6263–72. Available from: http://www.ncbi.nlm.nih.gov/pubmed/8887656

33. Avedis K, Eleanore AG, Grimes HL. The growth factor independence-1 transcription factor: New functions and new insights. Crit Rev Oncol Hematol [Internet]. 2006;59:85–97. Available from: http://linkinghub.elsevier.com/retrieve/pii/S1040842806000345

34. Jen J, Wang Y-C. Zinc finger proteins in cancer progression. J Biomed Sci [Internet]. 2016;23:53. Available from: http://jbiomedsci.biomedcentral.com/articles/10.1186/s12929-016-0269-9

35. Phelan JD, Shroyer NF, Cook T, Gebelein B, Grimes HL. Gfi1-cells and circuits: Unraveling transcriptional networks of development and disease. Curr. Opin. Hematol. 2010. page 300–7.

36. Vo BHT, Li C, Morgan MA, Theurillat I, Finkelstein D, Wright S, et al. Inactivation of Ezh2 Upregulates Gfi1 and Drives Aggressive Myc-Driven Group 3 Medulloblastoma. Cell Rep. 2017;18:2907–17.

37. Xing W, Xiao Y, Lu X, Zhu H, He X, Huang W, et al. GFI1 downregulation promotes inflammation-linked metastasis of colorectal cancer. Cell Death Differ. 2017;24:929–43.

38. Wassef M, Margueron R. The Multiple Facets of PRC2 Alterations in Cancers. J. Mol. Biol. 2017. page 1978–93.

39. Wang CG, Ye YJ, Yuan J, Liu FF, Zhang H, Wang S. EZH2 and STAT6 expression profiles are correlated with colorectal cancer stage and prognosis. World J Gastroenterol. 2010;16:2421–7.

40. Bachmann IM, Halvorsen OJ, Collett K, Stefansson IM, Straume O, Haukaas SA, et al. EZH2 expression is associated with high proliferation rate and aggressive tumor subgroups in cutaneous melanoma and cancers of the endometrium, prostate, and breast. J Clin Oncol. 2006;24:268–73.

41. Lo YH, Chung E, Li Z, Wan YW, Mahe MM, Chen MS, et al. Transcriptional Regulation by ATOH1 and its Target SPDEF in the Intestine. CMGH. 2017;3:51–71.

42. Bossuyt W, Kazanjian A, De Geest N, Van Kelst S, De Hertogh G, Geboes K, et al. Atonal homolog 1 is a tumor suppressor gene. PLoS Biol. 2009;7:0311–26.

43. Leopoldo S, Lorena B, Cinzia A, Gabriella DC, Angela Luciana B, Renato C, et al. Two subtypes of mucinous adenocarcinoma of the colorectum: Clinicopathological and genetic features. Ann Surg Oncol. 2008;15:1429–39.

44. Yang L, Cai Y, Zhang J, Hu H, Wenjing W, Wu Z, et al. Colorectal cancer with mucinous component compared to clinicopathological and molecular features as mucinous adenocarcinoma. J Clin Oncol [Internet]. 2017;35:e15166–e15166. Available from: http://ascopubs.org/doi/abs/10.1200/JCO.2017.35.15_suppl.e15166

45. Kazama Y, Watanabe T, Kanazawa T, Tada T, Tanaka J, Nagawa H. Mucinous carcinomas of the colon and rectum show higher rates of microsatellite instability and lower rates of chromosomal instability: A study matched for T classification and tumor location. Cancer. 2005;103:2023–9.

46. Oberg K, Castellano D. Current knowledge on diagnosis and staging of neuroendocrine tumors. Cancer Metastasis Rev. 2011;30:3–7.

47. Koh TJ, Dockray GJ, Varro A, Cahill RJ, Dangler CA, Fox JG, et al. Overexpression of glycine-extended gastrin in transgenic mice results in increased colonic proliferation. J Clin Invest. 1999;103:1119–26.

48. Korinek V, Barker N, Moerer P, Van Donselaar E, Huls G, Peters PJ, et al. Depletion of epithelial stem-cell compartments in the small intestine of mice lacking Tcf-4. Nat Genet. 1998;19:379–83.

49. Pinto D, Gregorieff A, Begthel H, Clevers H. Canonical Wnt signals are essential for homeostasis of the intestinal epithelium. Genes Dev. 2003;17:1709–13.

50. Wang Y, Giel-Moloney M, Rindi G, Leiter AB. Enteroendocrine precursors differentiate independently of Wnt and form serotonin expressing adenomas in response to active beta-catenin. Proc Natl Acad Sci [Internet]. 2007;104:11328–33. Available from: http://www.pnas.org/cgi/doi/10.1073/pnas.0702665104

